# Bayesian divergence-time estimation with genome-wide SNP data of sea catfishes (Ariidae) supports Miocene closure of the Panamanian Isthmus

**DOI:** 10.1101/102129

**Authors:** Madlen Stange, Marcelo R. Sánchez-Villagra, Walter Salzburger, Michael Matschiner

**Affiliations:** Department of Palaeontology and Museum, University of Zurich, Karl-Schmid-Strasse 4, 8006, Zurich, Switzerland; Zoological Institute, University of Basel, 4051 Basel, Switzerland; Centre for Ecological and Evolutionary Synthesis (CEES), Department of Biosciences, University of Oslo, 0316 Oslo, Norway

**Keywords:** Panamanian Isthmus, Central American Seaway, Bayesian inference, phylogeny, molecular clock, fossil record, SNPs, RAD sequencing, teleosts

## Abstract

The closure of the Isthmus of Panama has long been considered to be one of the best defined biogeographic calibration points for molecular divergence-time estimation. However, geological and biological evidence has recently cast doubt on the presumed timing of the initial isthmus closure around 3 Ma but has instead suggested the existence of temporary land bridges as early as the Middle or Late Miocene. The biological evidence supporting these earlier land bridges was based either on only few molecular markers or on concatenation of genome-wide sequence data, an approach that is known to result in potentially misleading branch lengths and divergence times, which could compromise the reliability of this evidence. To allow divergence-time estimation with genomic data using the more appropriate multi-species coalescent model, we here develop a new method combining the SNP-based Bayesian species-tree inference of the software SNAPP with a molecular clock model that can be calibrated with fossil or biogeographic constraints. We validate our approach with simulations and use our method to reanalyze genomic data of Neotropical army ants (Dorylinae) that previously supported divergence times of Central and South American populations before the isthmus closure around 3 Ma. Our reanalysis with the multi-species coalescent model shifts all of these divergence times to ages younger than 3 Ma, suggesting that the older estimates supporting the earlier existence of temporary land bridges were artifacts resulting at least partially from the use of concatenation. We then apply our method to a new RAD-sequencing data set of Neotropical sea catfishes (Ariidae) and calibrate their species tree with extensive information from the fossil record. We identify a series of divergences between groups of Caribbean and Pacific sea catfishes around 10 Ma, indicating that processes related to the emergence of the isthmus led to vicariant speciation already in the Late Miocene, millions of years before the final isthmus closure.

---

The emergence of the Isthmus of Panama had a profound impact on biodiversity in the Western Hemisphere. On land, the isthmus enabled terrestrial animals to migrate between the American continents, which led to massive range expansions and local extinctions during the so-called Great American Biotic Interchange (Woodburne 2010). In the sea, however, the rise of the isthmus created an impermeable barrier between the Caribbean and the Tropical Eastern Pacific (TEP), resulting in the geographic separation of formerly genetically connected marine populations (Lessios 2008). Due to its presumed simultaneous impact on speciation events in numerous terrestrial and marine lineages, the closure of the Isthmus of Panama has been considered one of the best biogeographic calibration points for molecular divergence-time estimation and has been used in several hundreds of phylogenetic studies (Bermingham et al. 1997; Lessios 2008; Bacon et al. 2015a). The precise age of isthmus closure assumed in these studies varies but generally lies between 3.5 Ma (e.g. Donaldson and Wilson Jr 1999) and 2.8 Ma (e.g. Betancur-R. et al. 2012), according to evidence from marine records of isotopes, salinity, and temperature, that all support an age in this range (Jackson and O’Dea 2013; Coates and Stallard 2013; O’Dea et al. 2016).

However, recent research has indicated that the history of the isthmus may be more complex than previously thought and that the isthmus may have closed temporarily millions of years before its final establishment around 3 Ma. The collision between the Panama Arc and the South American plate, which initiated the development of the isthmus, began as early as 25-23 Ma according to geochemical evidence (Farris et al. 2011). As a consequence, the Central American Seaway (CAS), the deep oceanic seaway connecting the West Atlantic and the East Pacific through the Atrato strait, is hypothesized to have narrowed down to a width of 200 km, still allowing for continued exchange between the oceans at this time (Farris et al. 2011; Montes et al. 2012). It has been argued that Eocene zircons in Colombian sediments support the existence of Miocene land bridges and fluvial connections between Panama and South America and thus a closure of the CAS around 15-13 Ma (Montes et al. 2015); however, alternative explanations for the occurrence of these zircons may be possible (O’Dea et al. 2016). Gradual shoaling of the CAS around 11-10 Ma has also been supported by biostratigraphic and paleobathymetric analyses (Coates et al. 2004) as well as seawater isotopic records (Sepulchre et al. 2014). On the other hand, a separate analysis of the seawater isotope records indicated that deep-water connections existed until around 7 Ma, followed by mostly uninterrupted shallow-water exhange (Osborne et al. 2014).

While the Atrato strait represented the main connection between the Caribbean and the Pacific throughout most of the Miocene, other passageways existed in the Panama Canal basin (the Panama isthmian strait) and across Nicaragua (the San Carlos strait) (Savin and Douglas 1985). Both of these passageways were likely closed around 8 Ma (and possibly earlier) but reopened around 6 Ma with a depth greater than 200 m, according to evidence from fossil foraminifera (Collins et al. 1996). The last connection between the Caribbean and the Pacific likely closed around 2.8 Ma (O’Dea et al. 2016), but short-lived breachings induced by sea-level fluctuations as late as 2.45 Ma cannot be excluded and receive some support from molecular data (Groeneveld et al. 2014; Hickerson et al. 2006).

In agreement with the putative existence of earlier land bridges, Miocene dispersal of terrestrial animals between North and South America is well documented in the fossil record. Fossils of a New World monkey, discovered in the Panama Canal basin, demonstrate that primates had arrived on the North American landmass before 20.9 Ma (Bloch et al. 2016). Furthermore, fossils of xenarthran mammals derived from South America (ground sloths, glyptodonts, and pampatheriids) were found in Late Miocene (9-8 Ma) deposits in Florida (Hirschfeld 1968; Laurito and Valerio 2012) and in Early Pliocene (4.8-4.7 Ma) deposits in Mexico (Carranza-Castañeda and Miller 2004; Flynn et al. 2005), and Argentinian fossils of the procyonid carnivore *Cyonasua* provide evidence that terrestrial mammals had also crossed from North to South America before 7 Ma (Marshall 1988; Bacon et al. 2016). The Argentinian fossils could still be predated by fossils of other mammalian North American immigrants in Late Miocene Amazonian deposits (Campbell et al. 2010; Frailey and Campbell 2012; Prothero et al. 2014); however, their age estimate of 9 Ma may require further confirmation (Carrillo et al. 2015). Dispersal of terrestrial animals is also supported by molecular data. Based on a metaanalysis of phylogenetic data sets, Bacon et al. (2015a,b) reported major increases in migration rates around 10-7 Ma and at 6-5 Ma. In combination with molecular evidence for increased vicariance of marine organisms around 10-9 Ma, the authors concluded that the Isthmus of Panama emerged millions of years earlier than commonly assumed.

Unfortunately, the observed evidence for dispersal of terrestrial organisms is in most cases insufficient for conclusions about the existence of earlier land bridges. This is due to the fact that most of these organisms are members of groups with a known capacity of oceanic dispersal (de Queiroz 2014), in many cases even over far greater distances than the gap remaining between North and South America in the Miocene (< 600 km; Farris et al. 2011). Before their dispersal to the North American landmass in the Early Miocene, primates had already crossed the Atlantic in the Eocene, when they arrived in South America (Kay 2015; Bloch et al. 2016). Many other mammal lineages have proven capable of oversea dispersal, which may be best illustrated by the rich mammalian fauna of Madagascar that is largely derived from Africa even though the two landmasses separated around 120 Ma (Ali and Huber 2010).

As a notable exception without the capacity of oversea dispersal, Winston et al. (2017) recently used Neotropical army ants (Dorylinae) to investigate the potential earlier existence of land bridges between North and South America. With wingless queens and workers that can only travel on dry ground, army ant colonies are highly unlikely to disperse across any larger water bodies (Winston et al. 2017) and are therefore particularly suited to answer this question. Based on restriction-site associated DNA sequencing (RAD-seq) and a concatenated alignment of genome-wide RAD-seq loci, Winston et al. (2017) generated a time-calibrated phylogeny that supported migration from South to Central America prior to 3 Ma for populations of the four species *Eciton burchellii* (4.3 Ma), *E. vagans* (5.5 Ma), *E. lucanoides* (6.4 Ma), and *E. mexicanum* (6.6 Ma). These estimates appear to support the existence of earlier land bridges; however, the results might be compromised by the fact that concatenation was used for phylogenetic inference. In the presence of incomplete lineage sorting, concatenation has not only been shown to be statistically inconsistent, with a tendency to inflate support values (Kubatko and Degnan 2007; Roch and Steel 2014; Linkem et al. 2016), but studies based on empirical as well as simulated data have also highlighted that concatenation may lead to branch-length bias and potentially misleading age estimates, particularly for younger divergence times (McCormack et al. 2011; Angelis and dos Reis 2015; Mendes and Hahn 2016; Meyer et al. 2017; Ogilvie et al. 2016, 2017).

A better alternative for more accurate estimates of divergence times related to the isthmus closure is the multi-species coalescent (MSC) model (Maddison 1997; Ogilvie et al. 2016, 2017). While the MSC also does not account for processes like introgression or gene duplication, it incorporates incomplete lineage sorting, which is likely the most prevalent cause of gene-tree heterogeneity in rapidly diverging lineages (Hobolth et al. 2007; Scally et al. 2012; Suh et al. 2015; Edwards et al. 2016). Unfortunately, available software implementing the MSC model either does not allow time calibration with absolute node-age constraints (Rannala and Yang 2003; Liu 2008; Kubatko et al. 2009; Liu et al. 2010; Bryant et al. 2012; Chifman and Kubatko 2014; Mirarab and Warnow 2015) or is computationally too demanding to be applied to genome-wide data (Heled and Drummond 2010; Ogilvie et al. 2017). To fill this gap in the available methodology, we here develop a new approach combining the Bayesian species-tree inference of the software SNAPP (Bryant et al. 2012) with a molecular clock model that can be calibrated with fossil or biogeographic constraints. SNAPP is well suited for analyses of genome-wide data as it infers the species tree directly from single-nucleotide polymorphisms (SNPs), through integration over all possible gene trees on the basis of the MSC model. By using SNPs as markers, SNAPP avoids the issue of within-locus recombination, a common model violation for almost all other implementations of the MSC (Lanier and Knowles 2012; Gatesy and Springer 2013, 2014; Springer and Gatesy 2016; Edwards et al. 2016; Scornavacca and Galtier 2017). SNAPP has been used in close to 100 studies (Supplementary Table S1), but with few exceptions, none of these studies inferred absolute divergence times. In five studies that estimated divergence times (Lischer et al. 2014; Demos et al. 2015; Ru et al. 2016; Portik et al. 2017; Cooper and Uy 2017), branch lengths were converted *a posteriori* to absolute times on the basis of an assumed mutation rate for the SNP set, a practice that should be taken with caution due to ascertainment bias (Lozier et al. 2016, also see the results of this study). With the possibility to analyze thousands of markers simultaneously, SNAPP nevertheless promises high precision in relative branch-length estimates, and accurate absolute divergence times when properly calibrated with fossil or biogeographic evidence.

We evaluate the accuracy and precision of our approach using an extensive set of simulations, and we compare it to divergence-time estimation based on concatenation. We then apply our method to reanalyze genomic data of Neotropical army ants with the MSC model, and we use it to estimate divergence times of Neotropical sea catfishes (Ariidae) based on newly generated RAD-seq data. Sea catfishes include species endemic to the TEP as well as Caribbean species in several genera. They inhabit coastal brackish and marine habitats down to a depth of around 30 m (Cervigón et al. 1993) and are restricted in dispersal by demersal lifestyle and male mouthbrooding. Sea catfishes are thus directly affected by geographic changes of the coast line, which makes them ideally suited to inform about vicariance processes related to the emergence of the Isthmus of Panama.

## Bayesian Divergence-Time Estimation with Simulated SNP Data

We designed five experiments based on simulated data to thoroughly test the performance of the MSC model implemented in SNAPP as a tool for divergence-time estimation with SNP data. In experiment 1, we tested the accuracy and precision of divergence times estimated with SNAPP and the degree to which these are influenced by the size of the SNP data set and the placement of node-age constraints. In experiment 2 we assessed the effect of larger population sizes, and in experiment 3 we tested how the precision of estimates depends on the number of individuals sampled per species. In experiment 4, we further evaluated SNAPP’s estimates of divergence times, the molecular clock rate, and the population size, based on data sets that include or exclude invariant sites, with or without ascertainment-bias correction. Finally, in experiment 5, we compared divergence-time estimates based on the MSC model implemented in SNAPP with those inferred with concatenated data using BEAST (Bouckaert et al. 2014). Characteristics of all simulated data sets are summarized in Table 1 Based on the results of experiments 1-5, we developed recommendations for divergence-time estimation with SNP data, and we then applied this approach to infer timelines of evolution for Neotropical army ants and sea catfishes.

**Table 1:**
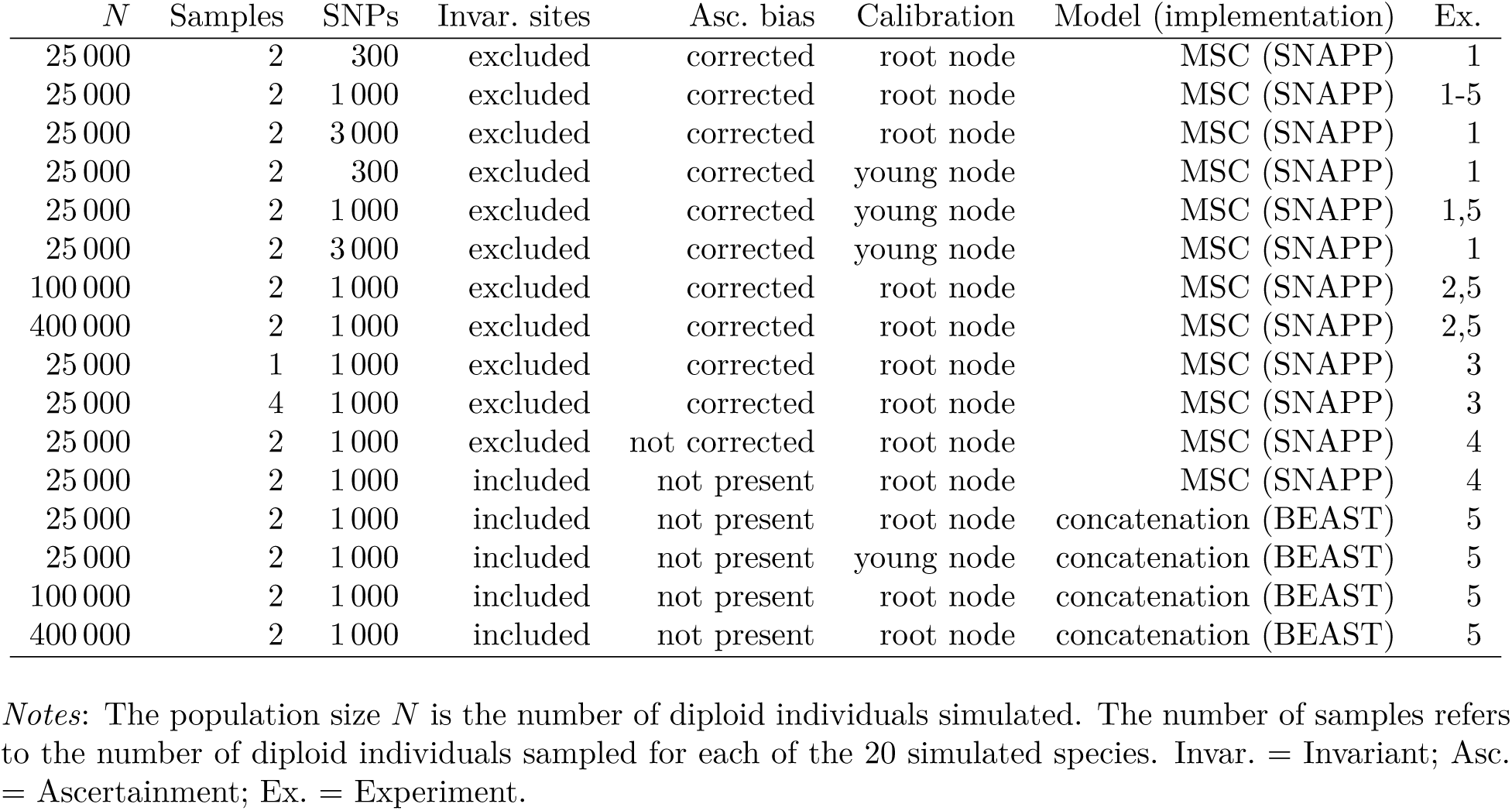
Simulated data sets and analysis settings used in experiments 1-5.

### Simulating Genome-Wide SNP Data

All simulation parameters, including the number of extant species, the age of the species tree, the population size, the generation time, the mutation rate, and the number of loci per data set were chosen to be roughly similar to those expected in empirical analyses with the software SNAPP (Supplementary Table S1). All simulated data sets were based on the same set of 100 species trees generated with the pure-birth Yule process (Yule 1925)(which is also the only tree prior currently available in SNAPP). Ultrametric species trees conditioned to have 20 extant species were generated with branch lengths in units of generations, using a constant speciation rate λ = 4 × 10^−7^ species/generation. Assuming a generation time of 5 years, this speciation rate translates to λ = 0.08 species/myr, within the range of speciation rates observed in rapidly radiating vertebrate clades (Alfaro et al. 2009; Rabosky et al. 2013). The ages of the resulting species trees ranged from 2.8 to 12.7 (mean: 6.5) million generations or from 14.2 and 63.6 (mean: 32.3) myr, again assuming the same generation time of 5 years.

For each simulated species tree, 10 000 gene trees were generated with the Python library DendroPy (Sukumaran and Holder 2010), using constant and equal population sizes for all branches. These population sizes were set to *N* = 25 000 diploid individuals for most analyses, but we also used the larger population sizes *N =* 100 000 and *N =* 400 000 in the simulations conducted for experiment 2 (Table 1). For each simulated gene tree, between 2, 4, or 8 terminal lineages were sampled per species, corresponding to 1, 2, or 4 diploid individuals per species (Table 1). Exemplary gene trees are shown in Supplementary Figure S1. Sequences with a length of 200 bp were then simulated along each of the gene trees with the software Seq-Gen (Rambaut and Grassly 1997), according to the Jukes-Cantor model of sequence evolution (Jukes and Cantor 1969) and a rate of 10^−9^ mutations per site per generation or 2 × 10^−4^ mutations per site per myr. The expected number of mutations per site between two individuals of a panmictic population, Θ, can be calculated as Θ = 4*Nμ*, where *N* is the number of diploid individuals, or half the number of haploid individuals, and *μ* is the mutation rate per site per generation. With the settings used in most of our simulations (*N* = 25 000; *μ =* 10^−9^), the expected number of mutations per site between two individuals of the same population is therefore Θ = 4 × 25 000 × 10^−9^ = 10^−4^.

At least 9 965 (mean: 9 998.5) of the resulting 10 000 alignments per species tree contained one or more variable sites. A single SNP was selected at random from all except completely invariable alignments to generate data sets of close to 10 000 unlinked SNPs for each of the 100 species trees. For each species, alleles of the 2, 4, or 8 terminal lineages sampled from each gene tree were combined randomly to form 1, 2, or 4 diploid individuals, which resulted in mean heterozygosities between 0.0012 and 0.0255. The resulting data sets of close to 10 000 unlinked SNPs were further subsampled randomly to generate sets of 300, 1000, and 3 000 bi-allelic SNPs for each species tree (see Table 1).

For the analyses in experiments 1-4, each of the 100 data sets of 300, 1 000, and 3 000 SNPs was translated into the format required for SNAPP, where heterozygous sites are coded with “1” and homozyguous sites are coded as “0” and “2”. Per site, the codes “0” and “2” were randomly assigned to one of the two alleles to ensure that the frequencies of these codes were nearly identical in each data set. For experiment 4 in which we tested for the effect of ascertainment bias in SNAPP analyses, the data sets of 1 000 SNPs were also modified by adding invariant sites. To each set of 1 000 SNPs, between 12184 and 32740 invariant sites (alternating “0” and “2”) were added so that the proportion of SNPs in these data sets matched the mean proportion of variable sites in the alignments initially generated for the respective species tree. Finally, for analyses using concatenation in experiment 5, we added the same numbers of invariant sites to the data sets of 1 000 SNPs; however, in this case we used the untranslated versions of these data sets with the original nucleotide code, and also used nucleotide code for the added invariant sites (randomly selecting “A”, “C”, “G”, or “T” at each site).

### Inferring Divergence Times from Simulated SNP Data

Input data and analysis settings were specified in the XML format used by SNAPP (Drummond and Bouckaert 2015); however, several important modifications were made to the standard analysis settings to allow divergence-time estimation with SNAPP. First, the forward and reverse mutation-rate parameters were both fixed to 1.0. By doing so, we assume a symmetric substitution model as well as equal frequencies, which is justified given that homozygous nucleotide alleles were translated into the codes “0” and “2” at random, independently at each site.

Second, we added a parameter for the rate of a strict molecular clock, the only clock model currently supported by SNAPP, and we used the one-on-x prior (Drummond and Bouckaert 2015) for the clock rate. Even though the one-on-x prior is improper (it does not integrate to unity), it is well-suited as a default rate prior because it combines the favorable attributes of i) giving preference to smaller values and ii) being invariant under scale transformations (Drummond et al. 2002). This means that regardless of the time scales spanned by the phylogeny of the investigated group, a relative change in the rate estimate (e.g. a multiplication by two) will always lead to the same relative change in the prior probability (e.g. a division in half). Thus, the one-on-x prior can be applied equally in analyses of groups with high or low mutation rates. However, it should be noted that because the one-on-x prior is improper, it is not suitable for model comparison based on estimates of the marginal likelihood. If such analyses were to be combined with divergence-time estimation in SNAPP, the one-on-x prior should be replaced with a suitable proper prior distribution.

The molecular clock rate was calibrated through age constraints on a single node of the species tree. To compare the effects of old and young calibrations, we conducted separate sets of analyses in which we placed this age constraint either on the root node or on the node with an age closest to one third of the root age (see Table 1). In each case, calibration nodes were constrained with log-normal calibration densities centered on the true node age. Specifically, these calibration densities were parameterized with an offset of half the true node age, a mean (in real space) of half the true node age, and a standard deviation of the log-transformed distribution of 0.1.

Third, initial tests indicated that our simulated data sets contained very little information about the ancestral population sizes on internal branches of the species tree, and that unreliable estimation of these population sizes could confound divergence-time estimates. We therefore decided not to estimate the population-size parameter Θ individually for each branch as is usually done in SNAPP analyses, but instead to estimate just a single value of Θ for all branches, assuming equal population sizes in all species. This assumption was met in our simulated data sets but may often be violated by empirical data sets; we turn to the implications of this violation in the Discussion. As a prior on Θ, we selected a uniform prior distribution, the only scale-invariant prior available for this parameter in SNAPP.

Finally, as we were interested in SNAPP’s ability to infer divergence times rather than the species-tree topology (which has been demonstrated previously; Bryant et al. 2012), we fixed the species-tree topology to the true topology. We provide a script written in Ruby, “snapp_prep.rb”, to generate XML input files for SNAPP corresponding to the settings described above (with or without a fixed species tree). Note that these settings, including the use of scale-invariant prior distributions, were deliberately not tailored towards our simulated data sets, but were instead intended to be generally applicable for divergence-time estimation with any SNP data set. As a result, the XML files produced by our script should be suitable for analysis without requiring further adjustments from the user. Our script is freely available at https://github.com/mmatschiner/snapp_prep. Details on operators used in our analyses are provided in Supplementary Text S1.

As SNAPP is specifically designed for the analysis of bi-allelic SNPs, its algorithm explicitly accounts for ascertainment bias introduced by the exclusion of invariable sites (Bryant et al. 2012; RoyChoudhury and Thompson 2012). Nevertheless, SNAPP allows invariant sites in the data set and the user may specify whether or not these have been excluded. Accordingly, we allow for invariant sites in all analyses of experiments 1-3, but not for the analyses of experiment 4 in which either ascertainment bias was not corrected for or invariant sites were added to data sets of 1 000 SNPs (see Table 1). This option did not apply to the analyses of concatenated data in experiment 5 as these were not conducted with SNAPP. As a substitution model, we applied the HKY model (Hasegawa et al. 1985) in analyses of concatenated data.

All XML files were analyzed using BEAST v.2.3.0 (Bouckaert et al. 2014) either with the SNAPP package v.1.3.0 (all analyses of experiments 1-4) or without additional packages (analyses of concatenated data sets in experiment 5). We performed between 400 000 and 18.4 million Markov-chain Monte Carlo (MCMC) iterations per SNAPP analysis and 500 000 iterations per concatenation analysis (Supplementary Table S2). Stationarity of MCMC chains was assessed by calculating effective samples sizes (ESS) for all parameters after discarding the first 10% of the chain as burn-in. Details on computational requirements of SNAPP analyses are given in Supplementary Text S2.

### Results: Precision and Accuracy of Parameter Estimates Based on Simulated SNPs

#### Experiment 1

In experiment 1, we tested the effects of data-set size and calibration placement on node-age estimates. A comparison of true and estimated node ages, for analyses of 100 data sets of 300, 1000, and 3 000 SNPs with node-age constraints on either the root or a younger node, is shown in Figure 1 and summarized in Table 2. As measured by the width of 95% highest posterior density (HPD) intervals, precision was generally greater for younger nodes and increased when larger numbers of SNPs were used for the analysis. In all sets of analyses, over 95% of the 95% HPD intervals contained the true age of the node, indicative of accurate inference free of node-age bias (Heath et al. 2014; Gavryushkina et al. 2014; Matschiner et al. 2017). The percentage of 95% HPD intervals containing the true node age was always slightly higher in analyses with root-node constraints even though the width of these HPD intervals was generally smaller.

**Table 2:** Accuracy and precision of node-age estimates (experiments 1-3).

| N | Samples | SNPs | Calibration | Accuracy (%) |  |  | Precision (myr) |  |  |  |
| --- | --- | --- | --- | --- | --- | --- | --- | --- | --- | --- |
|  |  |  |  | Young nodes | Old nodes | All | Young nodes | Old nodes | All | Ex. |
| 25 000 | 2 | 300 | root node | 95.3 | 97.2 | 96.1 | 3.89 | 7.73 | 5.46 | 1 |
| 25 000 | 2 | 1 000 | root node | 96.4 | 97.9 | 97.1 | 2.27 | 5.50 | 3.59 | 1-3 |
| 25 000 | 2 | 3 000 | root node | 96.4 | 99.6 | 97.7 | 1.50 | 4.47 | 2.71 | 1 |
| 25 000 | 2 | 300 | young node | 95.4 | 96.6 | 95.9 | 4.10 | 12.19 | 7.40 | 1 |
| 25 000 | 2 | 1 000 | young node | 96.2 | 97.2 | 96.6 | 2.44 | 7.85 | 4.65 | 1 |
| 25 000 | 2 | 3 000 | young node | 96.1 | 99.5 | 97.5 | 1.57 | 5.48 | 3.17 | 1 |
| 100 000 | 2 | 1 000 | root node | 95.2 | 98.3 | 96.5 | 2.28 | 5.67 | 3.66 | 2 |
| 400 000 | 2 | 1 000 | root node | 95.2 | 97.2 | 96.0 | 2.31 | 5.95 | 3.80 | 2 |
| 25 000 | 1 | 1 000 | root node | 95.9 | 98.7 | 97.1 | 2.28 | 5.46 | 3.58 | 3 |
| 25 000 | 4 | 1 000 | root node | 94.8 | 97.4 | 95.8 | 2.26 | 5.45 | 3.56 | 3 |

*Notes:* Accuracy was measured as the percentage of 95% HPD intervals containing the true node age. Precision was measured as the mean width of 95% HPD intervals for node-age estimates. Both measures are presented separately for young (true node age < 10 myr) and old (true node age > 10 myr) nodes. Ex. = Experiment.

**Figure 1:**
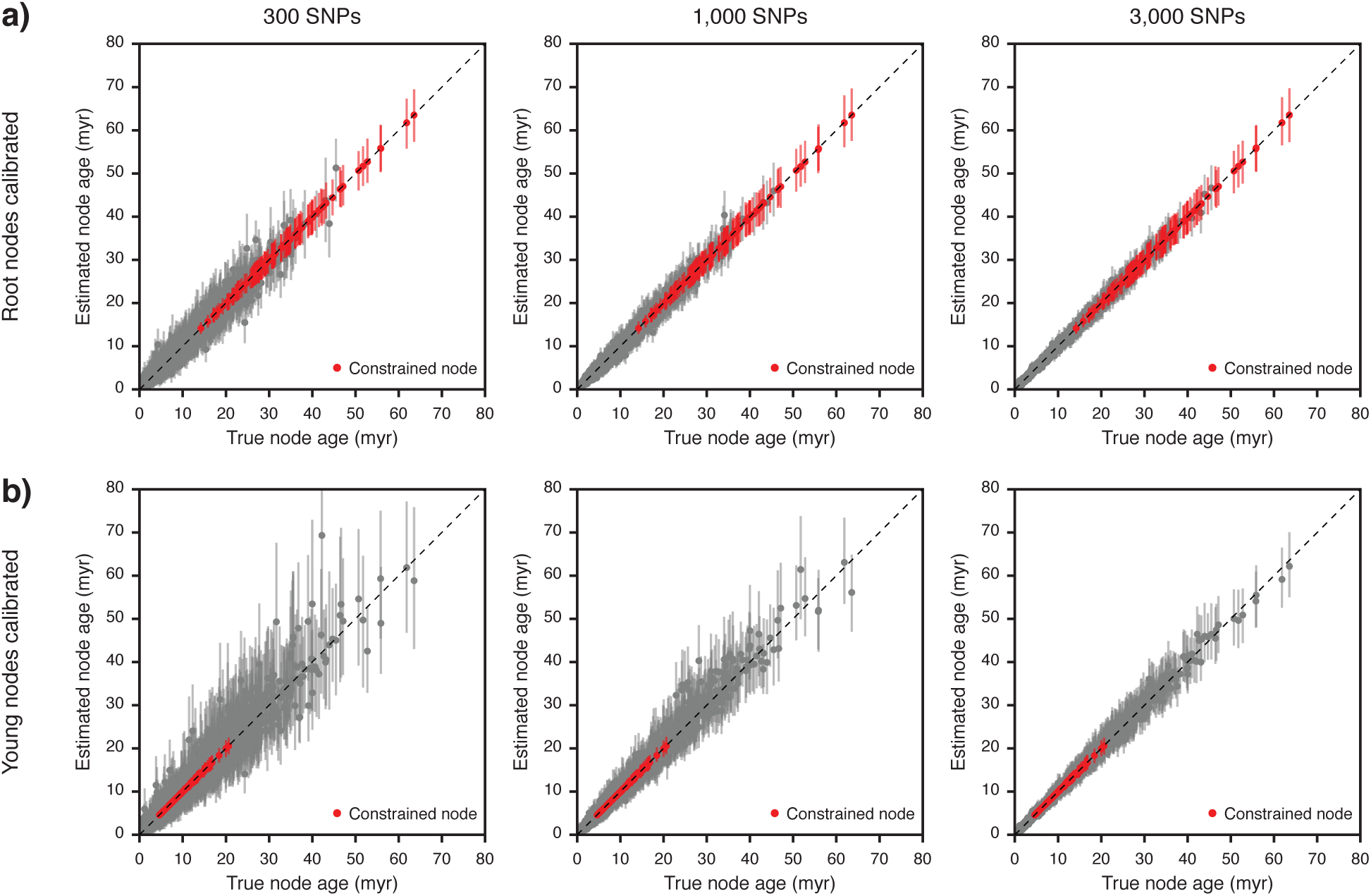
Comparison of true and estimated node ages (experiment 1). Results are based on 100 species trees and 300 to 3 000 SNPs generated per species tree. a) Node ages estimated with an age constraint on the root. b) Node ages estimated with an age constraint on a node that is approximately a third as old as the root. Mean age estimates of constrained and unconstrained nodes are marked with red and gray circles, respectively, and vertical bars indicate 95% HPD intervals.

#### Experiment 2

In experiment 2, we assessed the degree to which node-age estimates depend on the population sizes used in simulations. A comparison of node-age estimates obtained with simulated population sizes of *N* = 25 000, *N* = 100 000, and *N* = 400000 diploid individuals is shown in Supplementary Figure S2 and a summary of the accuracy and precision of these estimates is included in Table 2). The difference in the population sizes had only a negligible effect on node-age estimates: With all three population sizes, between 96.0 and 97.1% of 95% HPD intervals contained the true node age. The mean width of these intervals increased slightly from 3.59 myr with *N* = 25 000 to 3.80 myr with *N* = 400 000. While the difference in node-age estimates was minor, computational run times were significantly longer for analyses with larger population sizes (Supplementary Text S2, Supplementary Figure S3, and Supplementary Table S2).

#### Experiment 3

In experiment 3, we compared node-age estimates resulting from different numbers of individuals sampled from each species. The results for this comparison are shown in Supplementary Figure S4 and summary statistics are included in Table 2). With sample sizes of 1, 2, or 4 diploid indidivuals per species, the percentage of 95% HPD intervals containing the true node age remained between 95.8 and 97.1%, indicating accurate inference. The mean width of these intervals also remained mostly unchanged, between 3.56 and 3.59 myr (Table 2). In contrast, computational run times requrired for convergence were substantially longer with larger sample sizes: When a single diploid individual was sampled per species, MCMC analyses converged on average after 4.5 hours but required on average over 200 hours for convergence with a sample size of four individuals (Supplementary Text S2, Supplementary Figure S3, and Supplementary Table S2).

#### Experiment 4

In experiment 4, we tested SNAPP’s ability to recover the true clock rate, the true value of Θ, and the true population size when invariant sites were excluded from the data sets so that these consisted only of SNPs (as was the case for all data sets used in experiments 1-3). Regardless of whether SNAPP’s ascertainment-bias correction was used or not, the clock rates and Θ values estimated from data sets without invariant sites did not match the settings used for simulations (clock rate = 2 × 10^−4^ mutations per site per myr; Θ = 10^−4^; see above) (Fig. 2a,b, Table 3). While both parameters were underestimated roughly by a factor of three when ascertainment bias was corrected for, leaving this bias unaccounted led to parameter overestimation by more than an order of magnitude. Importantly, however, when ascertainment bias was accounted for, the resulting estimates of the population size *N* (calculated as *N* = Θ/4*μ* with *μ* being the mutation rate per generation, i.e., the estimated clock rate divided by the number of generations per myr) accurately recovered the true population size used for simulations (*N* = 25 000 in all simulations conducted for experiment 4; see Table 1), as 95% of the 95% HPD intervals included the true parameter value (Fig. 2c, Table 3). In contrast, the population size was underestimated when ascertainment bias was not corrected for: Mean estimates were on average 17.4% lower than the true population size and 35% of the 95% HPD intervals did not include the true parameter value (Fig. 2c, Table 3).

**Figure 2:**
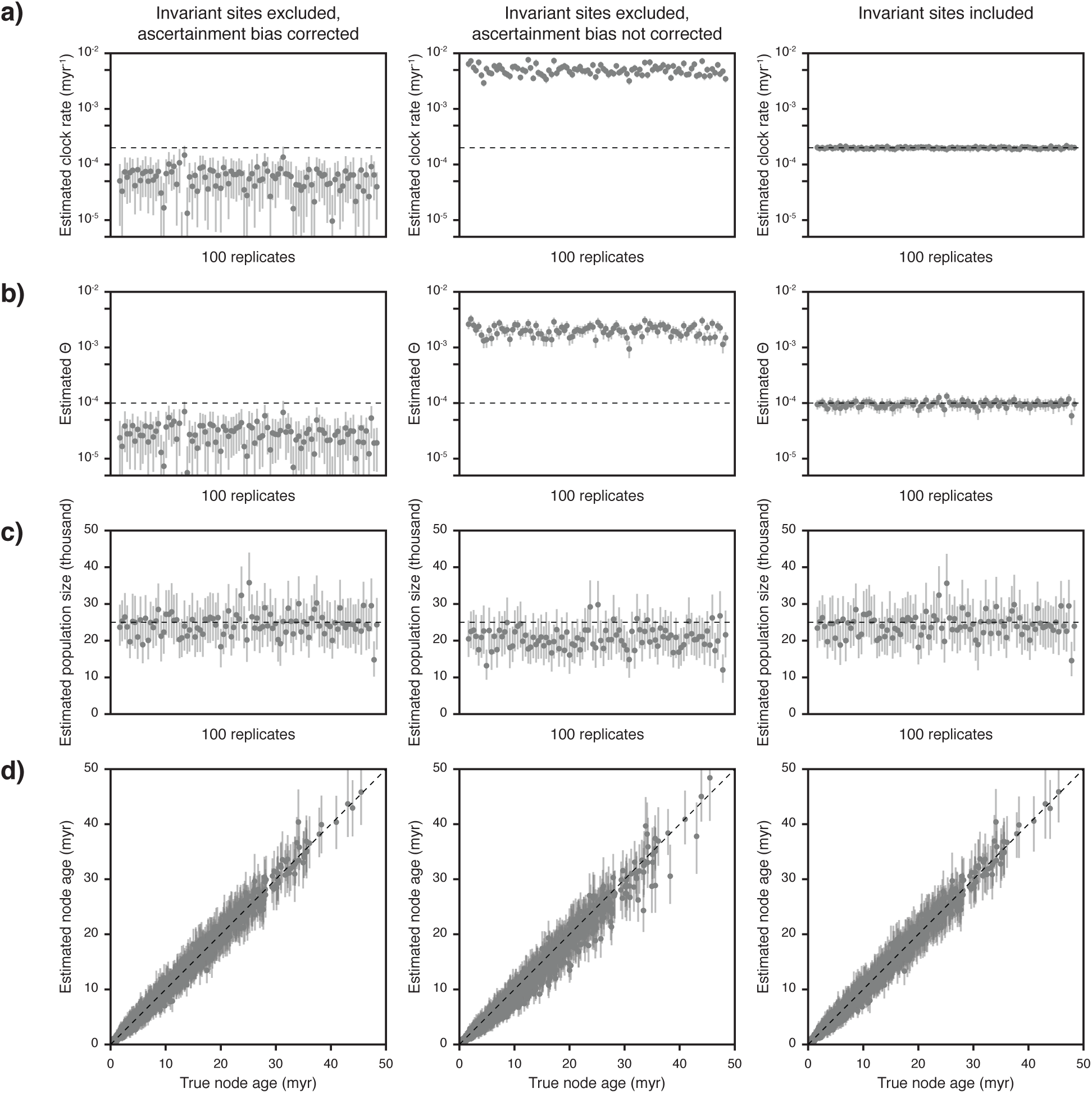
Estimates of node ages, the clock rate, Θ, and the population size, with and without ascertainment bias (experiment 4). Results are based on data sets of 1000 SNPs generated for each of 100 species trees, analyzed with and without SNAPP’s ascertainment-bias correction or after adding invariant sites to the data sets. Gray circles indicate mean estimates and 95% HPD intervals are marked with vertical bars. The visualization of node-age estimates in a) is equivalent to the illustration in Fig. 1, except that only unconstrained nodes are shown. Note that logarithmic scales are used for estimates of the clock rate (a) and Θ (b).

Our results of experiment 4 also showed that when invariant sites were excluded, SNAPP’s ascertainment-bias correction was required for the accurate estimation of node ages. Without ascertainment-bias correction, only 86.6% of the 95% HPD intervals contained the true node age (Fig. 2d, Table 3). Of the 13.4% of 95% HPD intervals that did not contain the true node age, almost all (13.2%) were younger than the true node age, indicating a tendency to underestimate node ages when ascertainment bias is not taken into account.

Instead of accounting for ascertainment bias, the inclusion of invariant sites also allowed the accurate estimation of the clock rate, the Θ-value, and the population size, with 100%, 90%, and 94% of the 95% HPD intervals containing the true values of these parameters, respectively (Fig. 2a-c, Table 3). Furthermore, the true node ages were also recovered reliably in these analyses and were included in 97.2% of the 95% HPD intervals (Fig. 2d).

**Table 3:** Estimates of clock rate, Θ, and the population size, in analyses of data sets with and without ascertainment bias (experiment 4).

| Inv. sites | Asc. bias | Mean estimate |  |  | Accuracy (%) |  |  |
| --- | --- | --- | --- | --- | --- | --- | --- |
| | | Clock rate | $\Theta$ | $N$ | Clock rate | $\Theta$ | $N$ |
| excluded | corrected | $5.93 \times 10^{-5}$ | $2.89 \times 10^{-5}$ | 24 438 | 2 | 2 | 95 |
| excluded | not corrected | $5.02 \times 10^{-3}$ | $2.05 \times 10^{-3}$ | 20 644 | 0 | 0 | 65 |
| included | not present | $1.99 \times 10^{-4}$ | $9.59 \times 10^{-5}$ | 24 226 | 100 | 90 | 94 |

*Notes:* The clock rate is given as mutations per site per myr. Inv. = Invariant; Asc. = Ascertainment.

**Figure 3:**
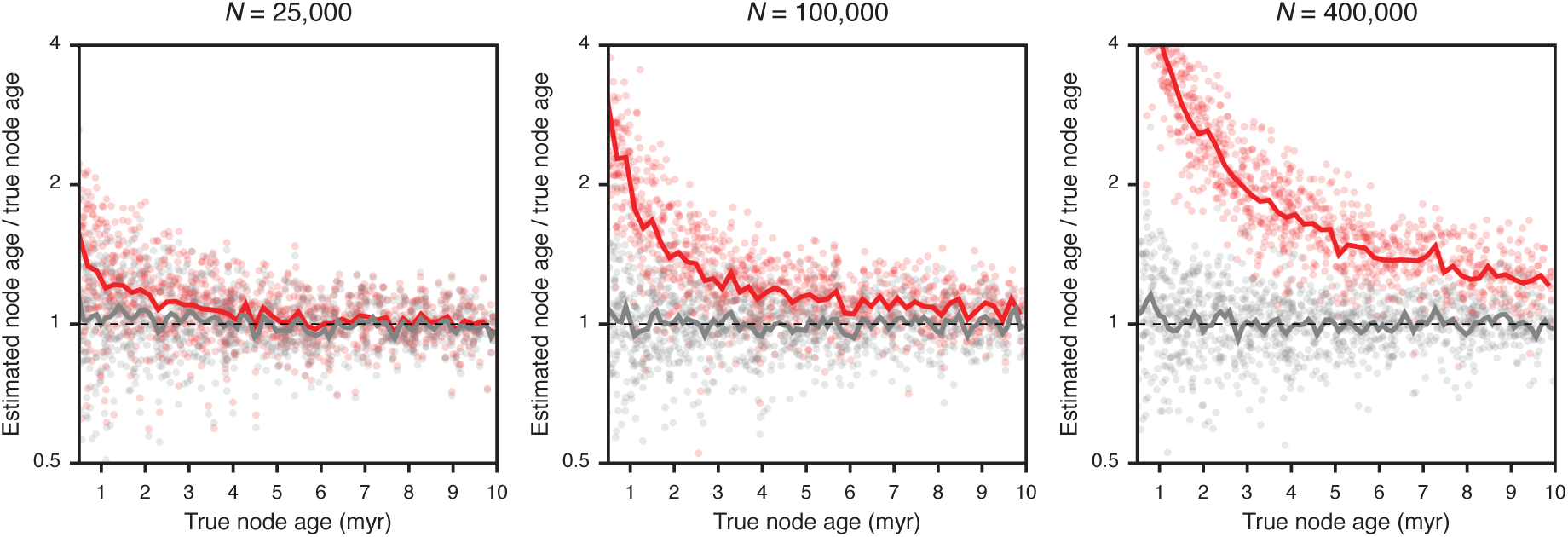
Error in node-age estimates obtained with the MSC or with concatenation (experiment 5). Results are based on analyses of 100 data sets of 1 000 SNPs, simulated with population sizes *N* = 25 000, *N* = 100 000, and *N* = 400 000. Gray and red dots indicate node-age estimates obtained with the MSC implemented in SNAPP and with BEAST analyses of concatenated data, respectively. Node-age error is measured as the ratio of the estimated node age over the true node age. Solid lines represent mean node-age errors in bins of 0.2 myr. Only nodes with true ages up to 10 myr are shown to highlight differences between the two methods. Note that a logarithmic scale is used for node-age error.

#### Experiment 5

In experiment 5, we compared node-age errors resulting from analyses with the MSC and with concatenation. This comparison indicated that both methods perform equally well for older nodes; however, the ages of younger nodes are commonly overestimated when concatenation is used. The degree of this overestimation increases with the population size: With a population size of *N* = 25 000, the ages of young nodes with a true node age between 0.5 myr and 10 myr were on average misestimated by 16.1% when using concatenation, but this percentage increased to 35.6% and 116.6% with the larger population sizes of *N* = 100 000 and *N* = 400 000, respectively (Fig. 3, Table 4). In contrast, the degree of misestimation of young node ages was not affected by population sizes when the MSC was used, and remained between 12.7% and 14.8%. With concatenation, the mean age estimate for a node with a true age around 3 Ma (±0.2 myr) increased from 3.3 Ma with *N* = 25 000 to 3.7 Ma with *N* = 100 000 and 5.9 Ma with *N* = 400 000, while mean age estimates for these nodes with the MSC remained between 3.0 and 3.2 Ma regardless of population size.

**Table 4:**
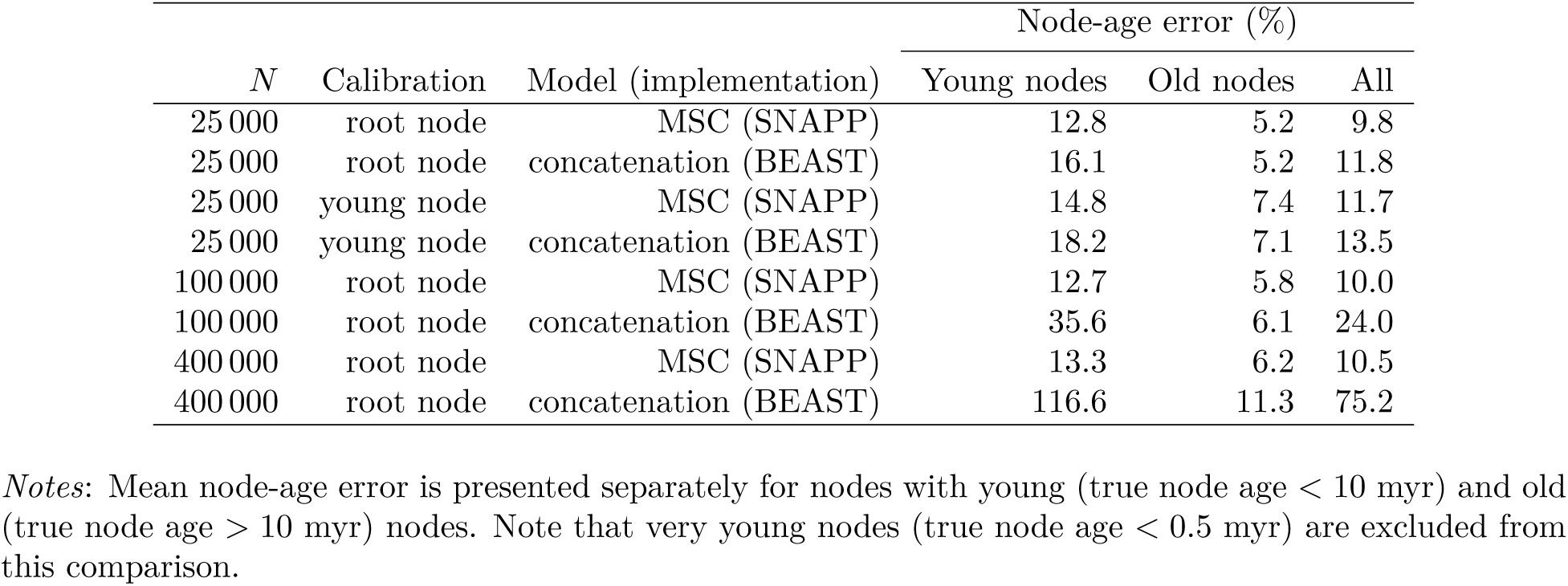
Mean error in node-age estimates in analyses using the MSC or concatenation, given in percent deviation from the true node age (experiment 5).

## Bayesian Divergence-Time Estimation with Empirical SNP Data

### Reanalysis of Neotropical Army Ant SNP Data

Divergence times of Neotropical army ants were estimated by Winston et al. (2017) based on a data set of 419 804 RAD-seq loci (39 927958 bp with 87.2% missing data), sequenced from 146 specimens of 18 species in five genera. Phylogenetic analysis of the concatenated data set led to divergence-time estimates older than 3 Ma between Central American and predominantly South American populations in each of four species of genus *Eciton (E. mexicanum, E. lucanoides, E. vagans,* and *E. burchellii*), which were taken as evidence for temporary land bridges prior to the full closure of the Panamanian Isthmus (Winston et al. 2017). To allow an efficient reanalysis of army ant divergence times with the MSC model, we reduced the size of this data set to the four specimens with the lowest proportions of missing data for each species, or for each of the two geographic groups in the four species *E. mexicanum, E. lucanoides, E. vagans,* and *E. burchellii.* We further filtered the data set so that maximally one SNP was included per RAD locus. The reduced data set included 413 bi-allelic SNPs suitable for analysis with SNAPP, with data available for at least one specimen per species. SNAPP input files in XML format were generated with the script “snapp prep.rb” (see above), using the same settings as for analyses of simulated data, except that the operator on the tree topology was not excluded. As in Winston et al. (2017), time calibration was based on the published age estimate of 37.23 Ma (confidence interval: 46.04-28.04 Ma) for the most recent common ancestor of Neotropical army ants (Brady et al. 2014). We specified this age constraint as a normally-distributed calibration density with a mean of 37.23 Ma and a standard deviation of 4.60 myr. To further reduce computational demands of the SNAPP analysis, we also enforced monophyly of each genus, and of each of the four species represented by two populations, according to the strong support (BPP: 1.0) that these groups received in Winston et al. (2017). We performed five replicate SNAPP analyses, each with a run length of 500 000 MCMC iterations. Chain convergence and stationarity were assessed through comparison of parameter traces among analysis replicates, using the software Tracer v.1.6 (Rambaut et al. 2014). As stationarity was supported by ESS values above 200 for all parameters in each analysis, MCMC chains of analysis replicates were combined after discarding the first 10% of each chain as burn-in. None of the ESS values of the combined chains were below 1 000, strongly supporting convergence of all analyses.

For comparison, we also repeated the analysis of army ant divergence times based on concatenation of all sequences, using a single specimen for each of the 22 species and geographic groups and excluding alignment positions with more than 50% missing data, which resulted in an alignment of 3 058 724 bp (with 37.1% missing data). Analyses based on concatenation were conducted in BEAST, using the GTR substitution model (Tavaré 1986) with gamma-distributed among-site rate variation and the same tree prior, clock model, and constraints as in analyses with the MSC. We again performed five analysis replicates, each with 600 000 MCMC iterations, and stationarity and convergence were again supported by ESS values above 200 in each individual analysis replicate and above 1 000 after combining the five MCMC chains.

### Results: Timeline of Neotropical Army Ant Diversification

Our reanalysis of Neotropical army ant SNP data with the MSC resulted in a strongly supported phylogeny (mean BPP: 0.94) that recovered the topology proposed by Winston et al. (2017) with the single exception that *Eciton mexicanum* appeared as the sister of *E. lucanoides* rather than diverging from the common ancestor of *E. lucanoides*, *E. burchellii*, *E. drepanophorum*, and *E. hamatum* (Supplementary Figure S5 and Supplementary Table S3). However, the timeline of army ant divergences inferred with the MSC was markedly different from the timeline estimated by Winston et al. (2017). Whereas Winston et al. (2017) estimated the crown divergence of the genus *Eciton* to have occurred around 14.1 Ma, our analysis based on the MSC placed this divergence around the Miocene-Pliocene boundary (5.48 Ma; 95% HPD: 7.52-3.52 Ma). In contrast to the previous analysis, the divergences between Central American and predominantly South American populations within *E. mexicanum* (1.82 Ma; 95% HPD: 3.02-0.76 Ma), *E. lucanoides* (2.47 Ma; 95% HPD: 3.88-1.22 Ma), *E. vagans* (0.33 Ma; 95% HPD: 0.71-0.05 Ma), and *E. burchellii* (0.54 Ma; 95% HPD: 1.12-0.13 Ma) were all placed in the Pleistocene in our study, in agreement with migration subsequent to the final isthmus closure. The population size inferred with the MSC, applying to all extant and ancestral species equally, was *N* = 53 854 (95% HPD: 34 433-75 294) diploid individuals, based on an assumed generation time of 3 years (Berghoff et al. 2008).

When using concatenation to estimate army ant divergence times, the mean age estimates of splits between Central American and predominantly South American lineages within *E. mexicanum* (2.47 Ma; 95% HPD: 3.09-1.88 Ma), *E. lucanoides* (3.74 Ma; 95% HPD: 4.68-2.83 Ma), *E. vagans* (1.31 Ma; 95% HPD: 1.65-1.00 Ma), and *E. burchellii* (2.07 Ma; 95% HPD: 2.56-1.55 Ma) were 35.7-397.1% older (Supplementary Figure S6 and Supplementary Table S4) than those based on the MSC model. While these age estimates for population splits in *E. mexicanum*, *E. vagans*, and *E. burchellii* would still agree with migration after the final closure of the isthmus, the confidence interval for the divergence time of populations within *E. lucanoides* does not include the accepted age for the final isthmus closure (2.8 Ma; O’Dea et al. 2016) and would thus support the existence of earlier land bridges.

### Generation of SNP Data for Neotropical Sea Catfishes

Twenty-six individuals that belong to 21 recognized species and two possibly cryptic species of the five Neotropical sea catfish genera *Ariopsis, Bagre, Cathorops, Notarius,* and *Sciades* were analyzed using RAD-seq (samples listed in Supplementary Table S5, including GPS coordinates and locality names). For four of these genera, our taxon set includes both species endemic to the TEP and species endemic to the Caribbean, hence, the divergences of these taxa were expected to have occurred prior to or simultaneously with the closure of the Panamanian Isthmus. Taxonomic identifications have previously been conducted for the same samples based on morphology as well as mitochondrial sequences (see Stange et al. 2016 for details) and were therefore considered to be reliable.

Fresh fin tissues were preserved in 96% ethanol for subsequent DNA extraction. DNA was extracted using the DNeasy Blood & Tissue Kit (Qiagen, Valencia, USA) following the manufacturer’s instructions. RNase treatment after digestion (but before precipitation) was performed in order to improve the purity of the samples. DNA concentrations were measured using a NanoDrop™ 1000 Spectrophotometer (Thermo Scientific, Waltham, MA, USA). The samples were standardized to 23.5 ng/μl and used to generate a RAD library, following the preparation steps described in Roesti et al. (2012) and using restriction enzyme Sbf1. We assumed a genome size of approximately 2.4 Gb as inferred from available C-values for sea catfishes (Gregory 2016). Therefore, we expected a recognition site frequency of 20 per Mb, which would yield around 50,000 restriction sites in total. Specimens were individually barcoded with 5-mer barcodes.

Two libraries were prepared and single-end sequenced with 201 cycles on the Illumina HiSeq 2500 platform, at the Department of Biosystems Science and Engineering, ETH Zurich. The resulting raw reads were demultiplexed (NCBI study accession: SRP086652) based on the individual barcodes with the script “process_radtags.pl” of the software Stacks v.1.32 (Catchen et al. 2011) and further analyzed with pyRAD v.3.0.5 (Eaton 2014). Settings of the pyRAD analysis included a minimum depth of 20 per within-sample cluster (Mindepth: 20), a maximum of four sites with a quality value below 20 (NQual: 4), maximally 20 variable sites within a cluster (Wclust: 0.89), and a minimum of 18 samples in a final locus (MinCov: 18). Quality filtering (step 2 in the pyRAD pipeline; Eaton 2014) resulted in the exclusion of 23-56% of the reads; after filtering, between 2.4 and 5.6 million reads remained per individual. Reads that passed the applied filtering steps resulted in about 40 000-166 000 within-sample clusters (step 3) with mean depths between 44 and 89. The estimated error rate and heterozygosity of these clusters (step 4) amounted to 0.0004-0.0009 and 0.0042-0.0107, respectively. Consensus-sequence creation from the within-sample clusters (step 5) based on the estimated heterozygosity and error rate, with a maximum of 20 variable sites, a minimal depth of 20, and additional paralog filtering (maximally 10% shared heterozygous sites), resulted in 21 575-38 182 consensus loci per sample. Between-sample clusters (step 6) were created with the same settings as within-sample clusters. These clusters were filtered again for potential paralogs (step 7) with a maximum of five shared heterozygous sites. The final data set contained 10 991-14064 clusters per individual. From these clusters, one SNP per locus was selected at random for use in phylogenetic inference, assuming that SNPs of different loci are effectively unlinked.

### Inferring the Divergence History of Neotropical Sea Catfishes

To incorporate existing estimates of the timeline of Neotropical sea catfish evolution into our analyses, we identified the age of the most recent common ancestor of the five sea catfish genera included in our taxon set *(Ariopsis, Bagre, Cathorops, Notarius,* and *Sciades)* from the time-calibrated phylogeny of Betancur-R. et al. (2012). Details of this phylogenetic analysis are given in Betancur-R. et al. (2012). In brief, Betancur-R. et al. (2012) used concatenation of five mitochondrial and three nuclear genes (a total of 7 190 sites) for phylogenetic inference of 144 species (representing 28 of the 29 valid genera of sea catfishes as well as diverse teleost outgroups), and divergence times were estimated with BEAST v.1.6.1 (Drummond et al. 2012) on the basis of 14 fossils and five biogeographic node-age constraints. However, as three of these biogeographic constraints were derived from an assumed closure of the Isthmus of Panama between 3.1 and 2.8 Ma and since our goal was to compare the timeline of Neotropical sea catfish evolution with the age estimates for the closure of the isthmus, we repeated the analysis of Betancur-R. et al. (2012) excluding these three constraints to avoid circular inference. All other analysis settings were identical to those used in Betancur-R. et al. (2012) but we used BEAST v.1.8.3, the latest version of BEAST compatible with the input file of Betancur-R. et al. (2012), and 150 million MCMC iterations for the inference.

The resulting age estimate for the most recent common ancestor of the genera *Ariopsis, Bagre, Cathorops, Notarius,* and *Sciades* (27.42 Ma; 95% HPD: 30.89-24.07 Ma) was then used as a constraint on the root of a species tree of Neotropical sea catfishes inferred with SNAPP, based on our RAD-seq data set of 21 sea catfish species. For this analysis, we used 1 768 bi-allelic SNPs for which data were available for at least one individual of each species or population. *Bagre pinnimaculatus* from Panama and *Sciades herzbergii* from Venezuela were represented by two individuals each, which were both considered as representatives of separate lineages in the SNAPP analyses. Differentiation between the populations from which these individuals were sampled was previously described based on morphology *(Bagre pinnimaculatus)* and distinct mitochondrial haplotypes (both species) (Stange et al. 2016). We again used our script “snapp prep.rb” (see above) to convert the SNP data set into SNAPP’s XML format.

The strict molecular clock rate was calibrated with a normally distributed calibration density (mean: 27.4182 Ma, standard deviation: 1.7 myr) on the root age, according to the result of our reanalysis of the Betancur-R. et al. (2012) data set. In addition, the fossil record of sea catfishes was used to define minimum ages for several lineages. The oldest fossil records of the genera *Bagre, Cathorops,* and *Notarius* have been described from the eastern Amazon Pirabas Formation on the basis of otolith and skull material (Aguilera et al. 2013). As the Pirabas Formation is of Aquitanian age (Aguilera et al. 2013), we constrained the divergences of each of the three genera with a minimum age of 20.4 Ma (Cohen et al. 2013). Furthermore, skull remains of the extant species *Sciades dowii, Sciades herzbergii, Bagre marinus,* and *Notarius quadriscutis* have been identified in the Late Miocene Urumaco Formation of northwestern Venezuela (Aguilera and de Aguilera 2004a), which therefore provides a minimum age of 5.3 Ma for these species. All fossils used for phylogenetic analyses are summarized in Supplementary Table S6.

We carried out five replicate SNAPP analyses, each with a run length of one million MCMC iterations, of which the first 10% were discarded as burn-in. Convergence was suggested by ESS values for all parameters above 200 in individual replicate analyses, and by ESS values above 1 000 after combining the output of the five replicates. The combined analysis output was used to sample a set of 1 000 trees as representative of the posterior tree distribution.

For the purpose of reconstructing ancestral distributions of sea catfishes taking into account the localities of fossil finds, eight fossil taxa were added to each of the 1 000 trees of the posterior tree set, according to their taxonomic assignment (Supplementary Table S6). For all additions, the age of the attachment point was chosen at random between the fossil’s age and the age of the branch to which the fossil was attached. We then used the posterior tree set including fossil taxa to infer the ancestral distribution of sea catfish lineages in the TEP or the Caribbean, based on stochastical mapping of discrete characters (Huelsenbeck et al. 2003) as implemented in function “make.simmap” of the phytools R package (Revell 2012). For this analysis, we assumed a uniform prior probability for the state of the root node and used an empirically determined rate matrix (Fig. 4, Supplementary Figure S7, and Supplementary Table S7). For comparison, we also performed a separate reconstruction of ancestral geography using the structured coalescent implementation of the BASTA package (De Maio et al. 2015) for BEAST. The reconstructed ancestral geographies were identical with both approaches; we therefore discuss only the results of the stochastic mapping approach below but provide results with both approaches in Supplementary Table S7.

### Results: Timeline of Neotropical Sea Catfish Diversification

The posterior distribution of species trees is illustrated in Figure 4 in the form of a cloudogram (Bouckaert and Heled 2014) with branches colored according to the stochastic mapping of geographic distribution. Our results suggest that the genus *Cathorops* is the outgroup to the other four genera (Bayesian posterior probability, BPP: 1.0) and that the earliest divergence between these groups probably occurred in what is now the Caribbean (BPP: 0.81). The four genera *Notarius, Bagre, Sciades,* and *Ariopsis* diverged (probably in this order; BPP: 0.92) in a rapid series of splitting events that occurred between 22 and 19 Ma, most likely also in the Caribbean (BPP: 0.89-1.0). Within-genus diversification of the sampled extant lineages began between 12 *(Notarius)* and 5 *(Ariopsis)* Ma, and these initial within-genus divergences occurred both within the Caribbean *(Sciades,* BPP: 1.0; *Bagre,* BPP: 0.77) and the TEP *(Ariopsis,* BPP: 0.89; *Cathorops,* BPP: 0.80). The most recent divergence between Caribbean and Pacific sea catfishes separated the Caribbean *Cathorops nuchalis* and *C. wayuu* from the Pacific *C. tuyra,* which occurred around 2.58 Ma (95% HPD: 3.37-1.87 Ma). Assuming a generation time of 2 years for sea catfishes (Betancur-R. et al. 2008; Meunier 2012), the estimated population size was *N* = 127250 (95% HPD: 105120-151900) diploid individuals.

**Figure 4:**
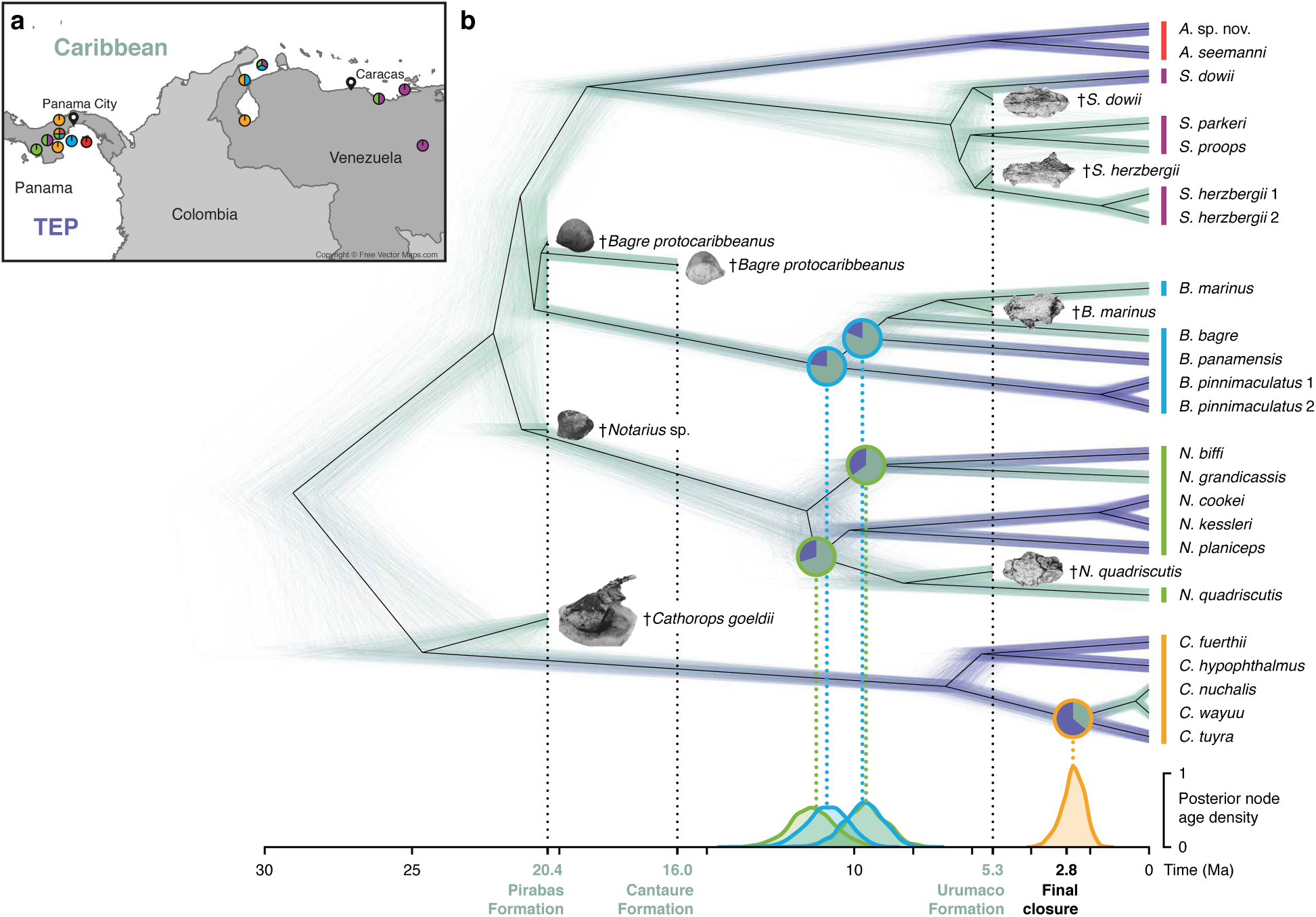
Time-calibrated species tree of Neotropical sea catfishes. a) Map of Panama and north-western South America with sampling locations of specimens used in this study. Colors of circles indicate genera of specimens sampled at a location: *Ariopsis,* red; *Sciades,* purple; *Bagre,* blue; *Notarius,* green; *Cathorops,* orange. b) Posterior distribution of time-calibrated species trees inferred with SNAPP, with fossil taxa added a posteriori (images of otoliths and partial skulls are from Aguilera and de Aguilera 2004b and from Aguilera et al. 2013, 2014; see Supplementary Table S6). Branch color indicates reconstructed geography: Caribbean; green, or Tropical Eastern Pacific (TEP); dark blue. Posterior densities of divergence times between Caribbean and Pacific lineages within *Notarius* (green), *Bagre* (blue), and *Cathorops* (orange) are shown below the species tree. Note that two divergence events around 10 Ma have nearly identical posterior density distributions: the divergence between *N. grandicassis* and *N. biffi* and the divergence between *B. panamensis* and the ancestor of *B. bagre* and *B. marinus*. Pie charts on nodes corresponding to divergences between Caribbean and Pacific lineages indicate posterior probabilities of ancestral distributions. All posterior estimates of node support, divergence times, and ancestral geography are summarized in Supplementary Table S7.

## Discussion

### Divergence-Time Estimation with Genome-Wide SNP Data

Our analyses based on simulated SNP data demonstrate that SNAPP, combined with a molecular clock model, allows the precise and unbiased estimation of divergence times in the presence of incomplete lineage sorting. As expected, the precision of estimates increased with the number of SNPs used for the analysis. With 3 000 SNPs, the largest number of simulated SNPs used in our analyses, uncertainty in divergence times resulted almost exclusively from the width of the calibration density (Fig. 1). In addition to data-set size, the placement of the node-age calibration also had an effect on the precision of divergence-time estimates, which was improved when the root node was calibrated instead of a younger node. This suggests that future studies employing divergence-time estimation with SNAPP should make use of constraints on the root node if these are available from the fossil record, from biogeographic scenarios, or from previously published time-calibrated phylogenies (as in our analyses of empirical SNP data of Neotropical army ants and sea catfishes). While we did not test the performance of multiple calibration points with simulated data, the use of additional calibration points can be expected to further improve the precision of divergence-time estimates; therefore these should be used if available.

It should be noted that even though all our analyses of both simulated and empirical data sets were calibrated through node-age constraints, this so-called “node dating” approach has been criticized for several reasons (Heath et al. 2014; O’Reilly et al. 2015; Matschiner et al. 2017). One problem associated with node dating is that prior distributions defined for node-age constraints are often chosen arbitrarily when minimum ages are provided by specific fossils but maximum ages are unknown. This problem has been adressed by using fossils as terminal taxa in “total-evidence dating” (Ronquist et al. 2012) and the “fossilized birth-death process” (Heath et al. 2014; Gavryushkina et al. 2017), but unfortunately, both of these approaches are not yet compatible with SNAPP. However, as a third alternative that overcomes the limitations of node dating, Matschiner et al. (2017) developed prior distributions for clade ages based on a model of diversification and fossil sampling and showed that these distributions allow unbiased inference when estimates for the rates of diversification and fossil sampling are available. The approach of Matschiner et al. (2017) is implemented in the CladeAge package for BEAST, which can be used in combination with SNAPP.

A limitation of our approach is the assumption of equal and constant population sizes on all branches of the phylogeny, which corresponded to the settings used in our simulations but may rarely be met in nature. Population growth or decline within a lineage is generally not estimated by SNAPP and may be only weakly identifiable in some cases Kuhner et al. (1998). Furthermore, the linking of population sizes was necessary to achieve feasible run times for analyses of data sets with around 20 species (with this number of species, assuming an individual population size for each branch would require an additional 37 model parameters). The single population-size parameter estimated with our method will therefore most commonly represent an intermediate value within the range of the true population sizes of the taxa included in the data set. As a result, divergence times might be slightly overestimated for groups in which the population size is underestimated and vice versa. Nevertheless, we expect that the degree of this misestimation is minor compared to the bias introduced by the alternative strategy of concatenation (Fig. 3, Table 4), which is equivalent to the MSC model only when all population sizes are so small that incomplete lineage sorting is absent and all gene trees are identical in topology and branch lengths (Edwards et al. 2016).

As a further limitation of our approach, only the strict molecular clock model is currently available in SNAPP; relaxed clock models such as the commonly used uncorrelated lognormal clock model of Drummond et al. (2006) have not yet been implemented. This means that particularly in clades that may be expected to have different mutation rates in different lineages, the precision of divergence-time estimates may be exaggerated, which should be considered in the interpretation of such results.

Our experiment 4 revealed that when SNP data sets are used without the addition of invariant sites, SNAPP’s estimates for the clock rate and Θ did not match those used in simulations (Fig. 2a,b, Table 3). While this mismatch might appear as a weakness of our approach, we do not consider it unexpected that these estimates change when slowly-evolving sites are excluded from the data set. Nevertheless, there are reasons why SNP-only data sets might be preferred over data sets that also include all invariant sites (Leaché and Oaks 2017). These reasons may be of practical nature, such as the comparative ease with which SNP-only data sets can be handled computationally due to their smaller file sizes, or the lower cost of genotyping when SNP arrays are used (even though these may by affected by additional biases; Leaché and Oaks 2017). A more important reason to use SNP-only data sets, however, is that determining whether or not sites are truly invariant is often not trivial due to low read coverage or mapping quality. As a result, the number of sites assumed to be invariant depends on the filters applied in variant calling and the ideal filtering settings that would result in the correct proportion of invariant sites are usually unknown. On the other hand, if the investigator chooses to focus exclusively on SNPs, strict filtering threshold can be applied that result in a conservative data set consisting only of sites that are known with high confidence to be variable. Based on the results of our analyses with simulated and empirical data, we argue that such data sets are highly suitable for phylogenetic inference with SNAPP, even though clock rate and Θ-values estimated from these data do not represent their genome-wide analogues. In our view, this mismatch is irrelevant for most phylogenetic analyses (even though users should be aware of it) because the clock rate and Θ usually represent nuisance parameters whereas the phylogeny, the divergence times, and the population size are of interest. As demonstrated in our experiments, all of these parameters of interest are estimated reliably from SNP data with our approach of divergence-time estimation with SNAPP, provided that SNAPP’s ascertainment-bias correction is applied.

### Insights Into the Taxonomy of Neotropical Sea Catfishes

Different views on the taxonomy of sea catfishes (Ariidae) have been supported by phylogenetic inference based on morphological features (Marceniuk et al. 2012b) and molecular data (Betancur-R. et al. 2007; Betancur-R. 2009). In the following, we address the most important differences between these views and how they are supported by our results, as well as new findings with regard to cryptic species.

#### Bagre and Cathorops

The morphology-based phylogenetic analysis of Marceniuk et al. (2012b) supported an earlier proposal by Schultz (1944) to raise the genus *Bagre* to family status due to its extraordinary morphological distinctiveness and its inferred position outside of a clade combining almost all other genera of sea catfishes. On the other hand, molecular studies have recovered *Bagre* in a nested position within sea catfishes, a position that is also supported by our results (Betancur-R. et al. 2007, 2012; Betancur-R. 2009). The proposed status of *Bagre* as a separate family is therefore not supported by molecular data. Instead of *Bagre,* our phylogeny identified the genus *Cathorops* as the sister of a clade combining *Notarius, Bagre, Sciades,* and *Ariopsis,* in contrast not only to morphology-based analyses but also to previous molecular studies that recovered a clade combining *Cathorops, Bagre,* and *Notarius,* albeit with low support (Betancur-R. et al. 2007, 2012; Betancur-R. 2009).

Within the genus *Bagre,* the existence of cryptic species has previously been suggested in *B. pinnimaculatus* based on cranio-morphological differences and distinct mitochondrial haplotypes of populations from the Bay of Panama and from Rio Estero Salado, Panama (Stange et al. 2016). Our current results corroborate this view, given that the estimated divergence time of the two populations *(B. pinnimaculatus* 1 and *B. pinnimaculatus* 2 in Fig. 5) is old (1.66 Ma; 95% HPD: 2.30-1.08 Ma) compared to the expected coalescence time within a species *(T_exp_* = 2 x *Ng* = 2 × 127 250 × 2 yr = 509 000 yr; with *N* according to SNAPP’s population size estimate and *g* according to an assumed generation time of two years for sea catfishes; Betancur-R. et al. 2008).

While *Cathorops nuchalis* has been declared a valid taxon based on morphological differentiation (Marceniuk et al. 2012a), mitochondrial sequences of this species were found to be indistinguishable from its sister species *C. wayuu* (Stange et al. 2016). In contrast, the nuclear SNP variation investigated here suggests that the two species are well differentiated and diverged 460 ka (95% HPD: 740-220 ka).

#### Notarius

According to our results, *Notarius quadriscutis* is either the sister to a Pacific clade composed of *N. cookei, N. kessleri,* and *N. planiceps* (BPP: 0.54), the sister to *N. biffi* and *N. grandicassis* (BPP: 0.07), or the sister to all other sampled extant members of the genus (BPP: 0.39). Based on morphology, the species has previously been placed in genus *Aspistor* together with *N. luniscutis* and the extinct *N. verumquadriscutis* (Marceniuk et al. 2012b; Aguilera and Marceniuk 2012). However, molecular phylogenies have commonly recovered species of the genus *Aspistor* as nested within *Notarius* (Betancur-R. and Acero P. 2004; Betancur-R. et al. 2012) and thus do not support the distinction of the two genera. Regardless of the exact relationships of *Notarius quadriscutis* in our species tree, our analyses suggest that the lineage originated around the time of the crown divergence of *Notarius* (11.61 Ma; 95% HPD: 13.23-10.21 Ma) and is thus younger than the earliest fossils assigned to the genus, *Notarius* sp. (Early Miocene; Aguilera et al. 2014). This implies that considering *Aspistor* as separate from *Notarius* would also require a reevaluation of fossils assigned to *Notarius.*

#### Ariopsis and Sciades

While molecular studies have supported the reciprocal monophyly of the genera *Ariopsis* and *Sciades* (Betancur-R. et al. 2007, 2012), species of the genus *Ariopsis* appeared paraphyletic in the morphology-based cladogram of Marceniuk et al. (2012b) and were there considered as members of *Sciades.* Our species tree inferred with SNAPP supports the results of previous molecular analyses since both genera appear as clearly monophyletic sister groups (BPP: 1.0) that diverged already in the Early Miocene (19.06 Ma; 95% HPD: 20.94-17.45 Ma).

Within *Sciades*, differentiation of mitochondrial haplotypes has been observed between brackish-water and marine populations of *S. herzbergii* from Clarines, Venezuela, and from the Golf of Venezuela (Stange et al. 2016). Our relatively old divergence-time estimate (1.64 Ma; 95% HPD: 2.20-1.04 Ma) provides further support for substantial differentiation of the two populations (S. *herzbergii* 1 and *S. herzbergii* 2 in Fig. 5) that could be driven by ecological adaptations to their contrasting habitats.

### Implications for the Closure of the Panamanian Isthmus

In agreement with our results based on simulated data, our reanalysis of genome-wide army ant data with both the MSC model and with concatenation indicated that recent divergence times can be overestimated if incomplete lineage sorting is not accounted for. As a result, the colonization of the North American landmass by army ants prior to the final closure of the Isthmus of Panama (2.8 Ma; O’Dea et al. 2016) was supported by our analyses using concatenation, but not by those using the MSC model. However, even the divergence times estimated with concatenation were generally younger than the divergence times reported by Winston et al. (2017), also on the basis of concatenation. This suggests that besides the variation introduced by the use of concatenation and the MSC, age estimates of army ant divergences were also influenced by other differences between our Bayesian divergence-time estimation and the analyses of Winston et al. (2017), which employed a penalized likelihood approach (Sanderson 2002) to estimate divergence times. These differences included not only the methodology used for time calibration, but also the number of specimens and alignment sites used in the analysis, as we had to filter the data set to comply with the assumption of the tree prior and to reduce the computational demands of the BEAST analysis. Nevertheless, our results suggest that previous claims of army ant migration to the North American landmass prior to the final isthmus closure (Winston et al. 2017) should be viewed with caution.

By combining Bayesian phylogenetic inference with reconstruction of ancestral geographic distributions, our analyses of sea catfish SNP data allowed us to estimate the timing and the location of divergence events separating lineages of Caribbean and Pacific sea catfishes (Fig. 5). The youngest of these events is the divergence of the Caribbean common ancestor of *Cathorops nuchalis* and *C. wayuu* from the Pacific *C. tuyra,* which we estimated to have occurred around 2.58 Ma (95% HPD: 3.37-1.87 Ma). As this age estimate coincides with the final closure of the Panamanian Isthmus around 2.8 Ma (O’Dea et al. 2016; Groeneveld et al. 2014), it appears likely that the closure was causal for vicariant divergence within *Cathorops.* According to our reconstruction of ancestral geographic distributions, the common ancestor of the three species *C. nuchalis, C. wayuu,* and *C. tuyra* more likely lived in the TEP (BPP: 0.64) than in the Caribbean. We note that this discrete type of inference may appear incompatible with the assumption that these lineages speciated through vicariance, given that in this case, the geographic distribution of the common ancestor should have extended across both regions as long as they were still connected. While our discrete ancestral reconstructions did not allow us to model this scenario of partially continuous distributions explicitly, our reconstructions can be reconciled with it if the inferred discrete geography is viewed not as the exclusive distribution of a species, but as the center of its distribution instead.

Surprisingly, the divergence of Caribbean and Pacific lineages within *Cathorops* was the only splitting event in our sample of sea catfishes that could be associated with the final closure of the Panamanian Isthmus around 2.8 Ma, even though the closure could be expected to affect a large number of species simultaneously. Instead, near-simultaneous divergence events between Caribbean and Pacific lineages were inferred at a much earlier time, about 10 Ma, in the genera *Bagre* and *Notarius.* Within *Notarius, N. grandicassis* of the Caribbean and the West Atlantic diverged from *N. biffi* of the TEP around 9.63 Ma (95% HPD: 10.99-8.30 Ma). This event may have coincided with the separation of Caribbean and Pacific lineages within *Bagre* (9.70 Ma; 95% HPD: 11.05-8.50 Ma), where the Pacific species *B. panamensis* diverged from a predominantly Caribbean (BPP: 0.81) ancestor that later gave rise to *B. bagre* and *B. marinus.* Two further divergence events between Caribbean and Pacific lineages of *Bagre* and *Notarius* were inferred slightly earlier, around 11 Ma. At 10.93 Ma (95% HPD: 12.29-9.60 Ma), the Pacific species *Bagre pinnimaculatus* diverged from the common ancestor of *B. marinus, B. bagre,* and *B. panamensis,* which likely had a distribution centered in the Caribbean (BPP: 0.77). Additionally, the common ancestor of the Pacific clade comprising *Notarius cookei*, *N. kessleri,* and *N. planiceps* diverged from the predominantly Caribbean (BPP: 0.70) lineage leading to *N. quadriscutis* at 11.29 Ma (95% HPD: 12.75-9.86 Ma).

Our time-calibrated species tree with reconstructed ancestral distributions (Fig. 5) shows further divergence events that separated Caribbean and Pacific lineages. The two sampled species of *Ariopsis* both occur in the TEP and diverged at about 19.06 Ma (95% HPD: 20.94-17.45 Ma) from the predominantly Caribbean genus *Sciades.* However, since *Ariopsis* also contains Caribbean species that we did not include in our data set, it remains unclear when and how often transitions between the Caribbean and the TEP took place in this genus. Caribbean origins of the genus *Cathorops* and of the species *Sciades dowii* are suggested by fossils from the Pirabas and Urumaco formations and indicate that these two lineages migrated to the Pacific after or simultaneous to the divergence from the fossil representatives. But since these divergence times were not estimated in our SNAPP analysis, the timing of migration of *Cathorops* and *Sciades dowii* also remains uncertain.

Regardless of these uncertainties, the near-simultaneous occurrence of several divergence events between Pacific and Caribbean lineages around 11-10 Ma suggests that geological processes associated with the emergence of the Panamanian Isthmus promoted vicariance long before the final closure of the isthmus around 2.8 Ma. Thus, even though our reanalysis of Neotropical army ant data suggested that army ants did not colonize the North American landmass before the final isthmus closure, our results based on sea catfish data add to the body of molecular evidence that indicates the emergence of temporary land bridges in the Late Miocene, leading to the separation of marine populations and migration of terrestrial animals (Donaldson and Wilson Jr 1999; Musilová et al. 2008; Bacon et al. 2015a,b; Carrillo et al. 2015; Acero P. et al. 2016; Huang 2016) long before the Great American Biotic Interchange (Woodburne 2010). While Miocene land bridges have been supported by a number of studies (Collins et al. 1996; Montes et al. 2015; Bacon et al. 2015a), it remains debated whether all of the connections between the Caribbean and the Pacific closed prior to 2.8 Ma, and whether they were blocked at the same time (O’Dea et al. 2016). Nevertheless, even if land bridges did not block all passages simultaneously, their emergence might have disrupted the distributions of catfish populations if these were localized in areas away from the remaining openings.

Although the rapid succession of divergence events between Caribbean and Pacific sea catfish lineages around 11-10 Ma indicates vicariance as the result of emerging land bridges, we cannot exclude that these events were driven by other modes of speciation, such as ecological speciation, and that their clustering within this relatively short period is coincidential. To discriminate between these possible explanations, a better understanding of the ecology of the diverging taxa will be important. In addition, the compilation of further diversification timelines for other groups of marine Neotropical species may strengthen the support for vicariance if divergences in these groups were found to cluster around the same times as in sea catfishes. As our results based on simulations suggest, these future analyses may benefit from genome-wide SNP data; however, concatenation should be avoided in favor of the MSC model to produce the most accurate estimates of divergence times. Importantly, our results clearly demonstrate that regardless of the causes of splitting events around 11-10 Ma, divergences between Caribbean and Pacific taxa are not necessarily linked to the final closure of the Panamanian Isthmus around 2.8 Ma.Thus, we reiterate earlier conclusions (Bacon et al. 2015a; De Baets et al. 2016) that the time of the final closure of the isthmus should no longer be used as a strict biogeographic calibration point for divergence-time estimation.

## Conclusion

We have demonstrated that the software SNAPP, combined with a molecular clock model, allows highly precise and accurate divergence-time estimation based on SNP data and the multi-species coalescent model. Our method thus provides molecular biologists with a powerful tool to investigate the timing of recent divergence events with genome-wide data. Our application of this method to two genomic data sets of Neotropical army ants and sea catfishes led to mixed support for the suggested closure of the Isthmus of Panama in the Miocene. We showed that army ants of the genus *Eciton* may have colonized the North American landmass only after the final closure of the Isthmus around 2.8 Ma and that previous conclusions supporting Miocene and Pliocene colonization events may have been influenced by branch-length bias resulting from concatenation. In contrast, we identify a series of four nearly coinciding divergence events around 10 Ma, as well as a final divergence around 2.8 Ma, between sea catfishes of the Caribbean and the TEP, which lends support to the hypothesis of Miocene isthmus closure and reopening. The rigorous application of divergence-time estimation with the multi-species coalescent model in future studies based on genomic data promises to contribute conclusive evidence for the timing and the effect of the emergence of the Panamanian Isthmus, one of the most significant events in recent geological history.

## Supplementary Material

Supplementary Material, including figures, tables, and input and output files of SNAPP and BEAST can be found in the Dryad Data Repository http://dx.doi.org/10.5061/dryad.f8k84. Code for all analyses is provided on https://github.com/mmatschiner/panama, and the script “snapp prep.rb” to generate SNAPP input files in XML format is available on https://github.com/mmatschiner/snappprep.

## Acknowledgements

We thank Remco Bouckaert for advice on divergence-time estimation with SNAPP and Emiliano Trucchi for testing our method. We are grateful to Richard Cooke for hospitality and assistance in coordinating our fieldwork in Panama, Aureliano Valencia, and Máximo Jiménez for help with sampling in Panama and Tito Barros, Cathy Villalba, and Jorge Domingo Carrillo-Briceño for field assistance and paperwork in Venezuela. We acknowledge help from Max Winston and Corrie Moreau to reuse their data set, and we thank Alexander Leow for contribution to SNP analyses. Part of this work was performed using the Abel high-performance computing facilies (www.hpc.uio.no) of the University of Oslo and the sciCORE (http://scicore.unibas.ch) scientific computing core facility at the University of Basel.

